# A hERG Blocker Facilitates K^+^ Channel Current by Agonizing Pore Opening while Blocking

**DOI:** 10.64898/2026.05.30.728541

**Authors:** Steffen S. Docken, Matthew J. Marquis, Khoa Ngo, Yuumu Wada, Satomi Kita, Vladimir Yarov-Yarovoy, Colleen E. Clancy, Igor Vorobyov, Timothy J. Lewis, Kazuharu Furutani, Jon T. Sack

**Author notes:** Contributed equally to this work (Steffen S. Docken, Matthew J. Marquis).

## Abstract

Many drugs that block voltage-gated K^+^ channels encoded by the human Ether-à-go-go-Related Gene (hERG) can cause long QT syndrome and life-threatening cardiac arrhythmias, yet the molecular mechanisms that determine this risk remain unclear. A process that may counteract arrhythmogenic hERG block, termed facilitation, is common to many clinically-approved hERG blockers including nifekalant, amiodarone, promethazine, imipramine, nortriptyline, haloperidol, verapamil, carvedilol, metoprolol, propranolol, quinidine, fluoxetine, and chlorpheniramine. Facilitation is an increase in hERG current, under certain conditions, due to these blockers.

Here, we propose that an agonism-while-blocking mechanism undergirds facilitation. We focus on nifekalant, a Class III antiarrhythmic drug and exemplar hERG blocker that induces facilitation. We tested the hypothesis that nifekalant opens hERG channel gates while blocking, and that unblock of these open-yet-blocked channels results in supranormal hERG current. We develop rate-theory kinetic models to identify features of agonism-while-blocking that produce facilitation. We generate atomistic models that predict that nifekalant blocks the hERG conduction path while modulating the intracellular conduction gate. Voltage-clamp measurements reveal that agonism-while-blocking undergirds nifekalant block and facilitation. We speculate that this agonism-while-blocking mechanism contributes to the relative safety of hERG blockers that induce facilitation.

## Introduction

The human Ether-à-go-go–Related Gene (*hERG*) encodes a voltage-gated potassium channel Kv11.1 pore-forming subunit. The mechanisms by which drugs affect hERG channels have important implications for human health. hERG channels underlie the rapid component of the cardiac delayed-rectifier potassium conductance that mediates the rapid recovery phase during the ventricular action potential (Sanguinetti *et al*., 1995; Sanguinetti and Tristani-Firouzi, 2006; Trudeau *et al*., 1995; Vandenberg *et al*., 2012). Drugs that decrease hERG conductance delay cardiac action potential termination and can result in long QT syndrome, potentially leading to fatal arrhythmias such as *torsade de pointes* (Kannankeril, Roden and Darbar, 2010; Roden, 2000; Roden, 2008; Sanguinetti *et al*., 1995; Sanguinetti and Tristani-Firouzi, 2006; Surawicz, 1989; Vandenberg *et al*., 2012). For this reason, advisory bodies warn against including hERG blockers in clinical trials (Gintant, Sager and Stockbridge, 2016; Leishman, Abernathy and Wang, 2020). However, hERG channels are unusually promiscuous targets for structurally diverse drugs, including many clinically important drugs, and designing drugs that do not inhibit hERG channels is a major challenge (Sanguinetti and Tristani-Firouzi, 2006; Vandenberg *et al*., 2012). As not all hERG blockers are proarrhythmic, wholesale exclusion of compounds with hERG-blocking activity results in the rejection of potentially valuable lead compounds from drug discovery pipelines (Gintant, Sager and Stockbridge, 2016). Indeed, numerous examples exist of drugs that block hERG and prolong the QT interval but have little proarrhythmic risk (De Ponti, Poluzzi and Montanaro, 2001; Redfern *et al*., 2003). Identifying distinguishing characteristics of safer hERG blockers could inform more nuanced assessment of lead compounds and save useful drugs from unwarranted rejection.

Curiously, many clinically approved hERG-blocking drugs produce a secondary agonistic effect on macroscopic hERG conductance called “facilitation” (Carmeliet, 1993; Furutani *et al*., 2019; Furutani *et al*., 2011; Hosaka *et al*., 2007; Jiang *et al*., 1999; Perry, Sanguinetti and Mitcheson, 2010; Yamakawa *et al*., 2012; Furutani, 2023). Here we dub hERG-blocking drugs that produce facilitation as “facilitatory blockers”, for brevity. Clinically approved facilitatory blockers include nifekalant (Igawa *et al*., 2002), amiodarone, promethazine, imipramine, nortriptyline, haloperidol, verapamil, carvedilol, metoprolol, propranolol, quinidine, fluoxetine, and chlorpheniramine (USFDA, 2025; Furutani *et al*., 2011; Yamakawa *et al*., 2012). Facilitation is proposed to increase hERG conductance in ventricular myocytes during the recovery phase of action potentials and thereby may lower the risk that a *hERG* blocker will cause arrhythmia (Furutani *et al*., 2019). However, the molecular mechanism of facilitation remains unclear. The widespread clinical use of facilitatory hERG blockers suggests that elucidating the molecular mechanism of facilitation could reveal predictors of clinical safety for hERG blockers.

The conformational changes of hERG channel gating interact with facilitatory blockers. hERG is a tetramer with each subunit containing 6 membrane-spanning helices, designated S1-S6 (Asai *et al*., 2021; Robertson and Morais-Cabral, 2020; Wang and MacKinnon, 2017). At the intracellular end of the channel pore, the S6 helices can constrict, creating a voltage-activated gate for potassium and drug entry into the membrane-buried central cavity of the channel (Robertson and Morais-Cabral, 2020; Vandenberg *et al*., 2004). Without closing the S6 gate, open hERG channels can transition to an inactivated state (Vandenberg *et al*., 2004) via relatively rapid, voltage-dependent conformational changes in the selectivity filter located at the extracellular end of the central cavity (Li *et al*., 2021; Trudeau *et al*., 1995). Inactivation markedly reduces hERG conductance at voltages that elicit maximal S6 gate opening, such as in the plateau phase of the cardiac action potential. Upon repolarization, hERG channels close their S6 gate slowly, but rapidly recover from inactivation. During repolarization of cardiac action potentials, recovery from inactivation increases hERG current, accelerating repolarization. When hERG is blocked, loss of its repolarizing current can trigger cardiac arrhythmias.

Facilitation, like block, is proposed to occur due to drug binding in the hERG central cavity lined by S6 helices (Hosaka *et al*., 2007). However, the structural and kinetic differences between facilitation and block remain to be elucidated. Facilitation is characterized by a negative shift in the voltage eliciting half-maximal hERG tail conductance (activation *V*_1/2_) under certain conditions (Furutani *et al*., 2011; Perry, Sanguinetti and Mitcheson, 2010). Due to this shift, some voltage steps can elicit more current than in the absence of facilitatory blockers (Figure 1) (Furutani *et al*., 2019; Furutani *et al*., 2011; Hosaka *et al*., 2007; Yamakawa *et al*., 2012). In addition to the presence of a facilitatory blocker, facilitation requires prior voltage stimulation.

**Figure 1.**
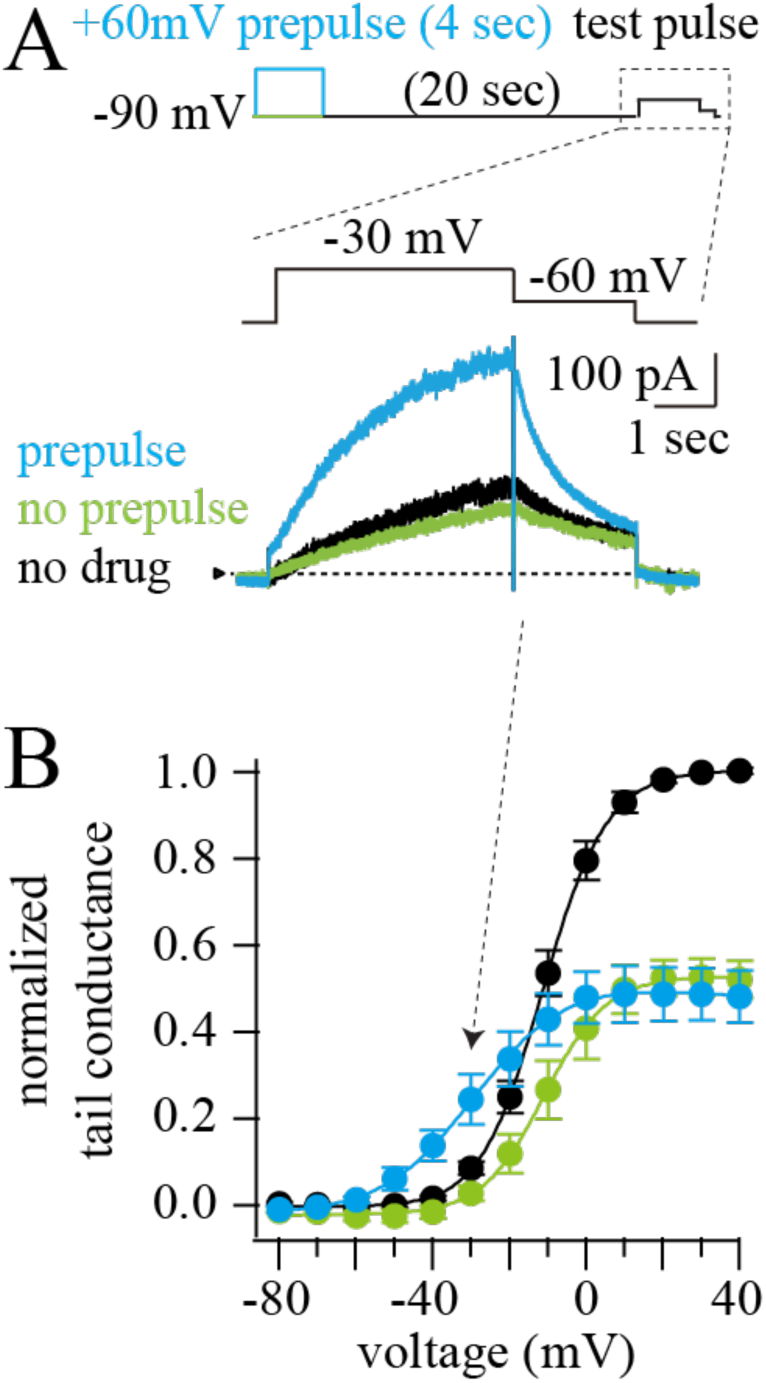
The pore blocker nifekalant facilitates hERG currents. (**A**) hERG currents in response to voltage-clamp pulses under control condition after 4s +60 mV prepulse (black), in 100 nM nifekalant with prepulse (blue), in 100 nM nifekalant without prepulse (green). (**B**) The relationship between the tail current amplitude and test pulse voltage. Means ± SEM (*n* = 5 cells). Data replotted from Furutani *et al*., 2019. Prepulse does not change hERG conductance without nifekalant (Furutani *et al.,* 2011).

Facilitation can be induced by a ‘pre-pulse’ to a voltage more positive than the activation *V*_1/2_ lasting hundreds of milliseconds or longer (Carmeliet, 1993; Furutani *et al*., 2011; Hosaka *et al*., 2007; Jiang *et al*., 1999; Yamakawa *et al*., 2012). Without a pre-pulse, facilitatory blockers inhibit hERG conduction at all voltages. We define facilitation as an increase in hERG conductance below the conductance-voltage (G–V) midpoint, following a depolarizing pre-pulse.

Facilitation has been studied in-depth with the Class III antiarrhythmic agent nifekalant. In the presence of nifekalant, channel opening initiates block, and recovery occurs at voltages that promote channel closing (Furutani *et al*., 2022; Kushida *et al*., 2002). This suggests that open or inactivated channels have greater nifekalant affinity than closed channels. However, increased inactivation does not increase block by nifekalant (Kushida *et al*., 2002), suggesting that S6 gate opening, not inactivation, increases nifekalant blocking affinity. In the presence of nifekalant, facilitation induced by a pre-pulse produces a shift of approximately –25 mV in the activation *V*_1/2_ (Furutani *et al*., 2011; Hosaka *et al*., 2007) (Figure 1B). The requirement that nifekalant must enter through the open channel pore explains the voltage dependence of entry into the facilitated state (Furutani *et al*., 2022). Mutation of cavity-lining residues can affect block and facilitation by nifekalant differently (Hosaka *et al*. 2007), suggesting that distinct binding poses underlie facilitation. Since facilitatory blockers enhance conductance, nifekalant might be expected to bind in a manner that does not block the flow of potassium through the channel but, rather, stabilizes a conductive state of the channel. If such a non-blocking binding mode exists, potassium ions would need to permeate the central cavity while nifekalant is bound. However, structures of open hERG channels determined by cryo-electron microscopy revealed a notably narrow cavity (Asai *et al*., 2021; Wang and MacKinnon, 2017), prompting us to question whether it is plausible for potassium ions to traverse a cavity occupied by nifekalant.

Here, we propose a mechanism of facilitation in which the facilitatory compound always blocks the pore while bound to the channel. We call this mechanism “agonism-while-blocking” because its key feature is that facilitatory blockers agonize S6 gate opening while blocking the central cavity. Agonism-while-blocking is a simpler alternative to the proposal that facilitatory blockers have a second binding site that opens channels. Here we address whether agonism-while-blocking could explain the behavior of nifekalant.

Using a simple kinetic rate theory model, we demonstrate that agonism-while-blocking can produce facilitation. We then analyze atomistic models of nifekalant bound in the cavity of open and putative closed models of the hERG channel to assess whether structural features of these complexes are consistent with agonism-while-blocking. Using whole-cell voltage clamp of hERG1a expressed in HEK293 cells, we investigate whether hyperpolarizing voltage can trap channels in the more readily opened ’facilitated’ state, just as it traps nifekalant in its blocking site. We then measure the unblocking kinetics of open channels and conformational state-dependent affinity changes, to develop the agonism-while-blocking mechanism into a kinetic rate theory model of hERG facilitation.

## Materials and Methods

### Cell preparation and hERG channel current recording

A human embryonic kidney (HEK) 293 cell line stably expressing hERG was kindly provided by Dr. Craig T. January and maintained in minimum essential medium supplemented with 10% fetal bovine serum (FBS; biowest, Nuaillé, France) and 400 µg/ml G418 (InvivoGen, San Diego, CA) as previously described (Zhou *et al*., 1998). HEK cells used for electrophysiological study were adhered to poly-L-lysine-coated (MW 30,000-70,000, Sigma-Aldrich; 0.1 mg/mL for 2 hr at 37 °C) coverslips (CS-5R 5 mm diameter coverslips, Warner Instruments Corp., MA, USA) in 12-well plates and transferred to a small recording chamber mounted on the stage of an inverted microscope (Olympus IX71, Olympus, Tokyo, Japan), and were continuously superfused with HEPES-buffered Tyrode solution containing (in mM) 137 NaCl, 4 KCl, 1.8 CaCl_2_, 1 MgCl_2_, 10 glucose, and 10 HEPES (pH 7.4 with NaOH). Membrane currents were recorded in a whole-cell configuration established using pipette suction (Hamill *et al*., 1981). Leak compensation was not used. The borosilicate micropipettes (G150TF-4, Warner Instruments Corp., MA, USA) had a resistance of 2–3 MΩ when filled with the internal pipette solution containing (in mM) 120 KCl, 5.374 CaCl_2_, 1.75 MgCl_2_, 10 EGTA, 10 HEPES (pH 7.2 with KOH). Liquid junction potential with this internal solution was calculated as smaller than −4 mV (Liquid Junction Potential Calculator, https://swharden.com/software/LJPcalc/app/), and the offset was not corrected. Series resistance was typically under 5 MΩ. Series resistance compensation was used when needed to constrain voltage error to <10 mV. Holding potential was -90 mV. Whole-cell recordings were performed using an Axopatch 200B patch-clamp amplifier (Molecular Devices, Sunnyvale, CA), ITC-18 interface and PatchMaster software (HEKA Elektronik, Lambrecht, Germany) or Dendrite interface and SutterPatch software (Sutter Instrument, Novato, CA). The data were stored on a computer hard disk and analyzed using PatchMaster and Igor Pro 9 (WaveMetrics, Portland, OR). Nifekalant was obtained from Cayman Chemical (Ann Arbor, MI).

### hERG docking simulations

Models of nifekalant docked to open and closed hERG channels were generated as described, and the data presented here are original analyses of structural models in a dataset also analyzed previously (Ngo *et al*., 2025). The top 100 docking poses were clustered by ligand root-mean-square deviation (RMSD). To account for the homotetrameric symmetry of the hERG channel, the RMSD calculation compared each ligand pose against all other poses, including their equivalents rotated by 90°, 180°, and 270° around the central pore axis. Using an RMSD threshold of 2.5 Å, clusters containing at least 8 poses were considered significant; those with fewer were deemed outliers. The top three clusters with the most favorable average binding free energies had their top scoring poses chosen for further analysis. These representative poses underwent a detailed examination of protein-ligand interactions using the Grapheme Toolkit from the OpenEye software suite (https://www.eyesopen.com), with interaction detection criteria modified as specified in the OEChem Toolkit manual (https://docs.eyesopen.com/toolkits/python/oechemtk/OEBioClasses/OEPerceiveInteractionOpti ons.html). The interaction patterns and binding sites were visualized as two-dimensional images and detailed spatial arrangements were explored through three-dimensional visualizations using ChimeraX software (Pettersen *et al*., 2021).

### Mathematical models of drug-hERG interaction

The minimal drug-*hERG* model consists of a closed state (𝐶), an open state (O), a drug-bound closed state (𝐶_*_), and a drug-bound activated (O*) state. Activation and deactivation rate constants are defined as 𝛼 and 𝛽 for non-drug-bound channels and *f*_𝛼_𝛼 and *f*_𝛽_𝛽 for drug-bound channels. Drug can only bind to the O state and unbind from the O* state with the rate constants [𝐷]𝑘_block_ and 𝑘_unblock_, respectively. The full set of differential equations describing the model dynamics are

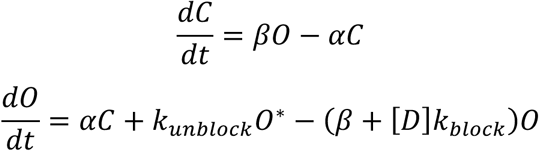

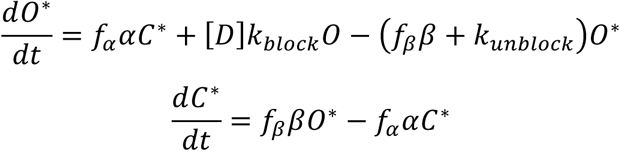

The rate constants 𝛼(𝑉) and β(𝑉) are modeled as single exponentials with the forms

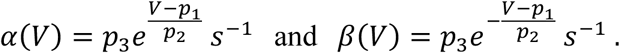

The parameters *p_1_*, *p_2_*, and *p_3_*, were set by first performing a least squares fit of 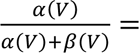 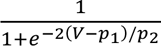 to the steady state activation data from our experiments (Figure 2B). Then, *p_3_* was used to scale the time constant of activation, 1/(𝛼(𝑉)+𝛽(𝑉)), to 1 s at the relevant test pulse potential of −40 𝑚𝑉 (used in Figure 2), which is consistent with the value observed experimentally (Figure 4F). The parameters are 𝑝_1_ = −11.0 𝑚𝑉, 𝑝_2_ = 16.2 𝑚𝑉, and 𝑝_3_ = 0.162. The dissociation constant (*K_D_*) plotted in Figure 2G is defined as the concentration of drug that leads to 50% of channels being bound to drug at steady state and is given by the equation

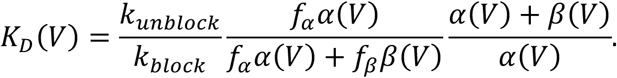

**Figure 2.**
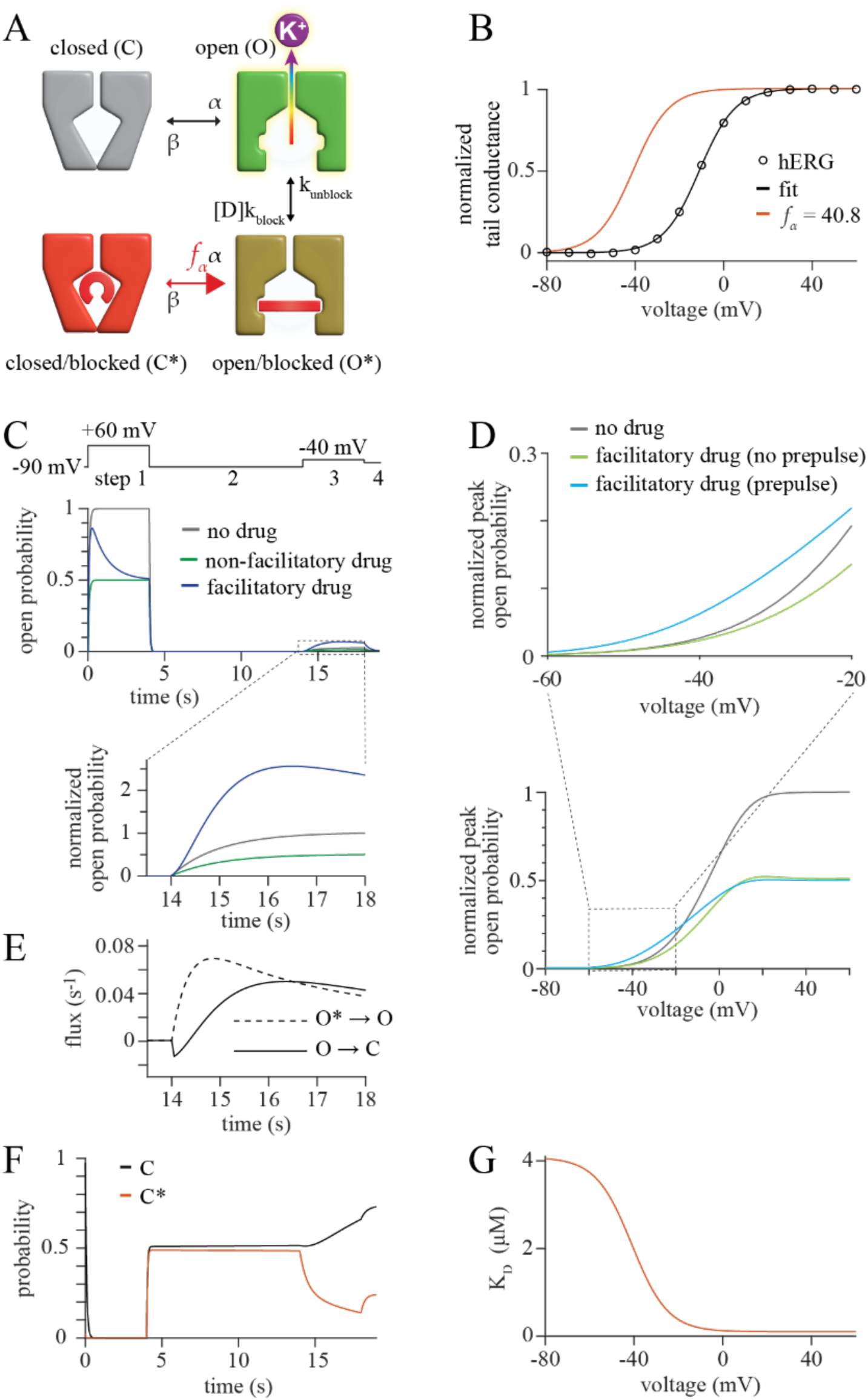
Agonism-while-blocking can produce facilitation. (**A**) Cartoon depicting the proposed state transitions underlying facilitation. Closed channels (State C, gray) can open (State O, green) and open channels can become blocked (State O*, yellow). Open/blocked channels can close (State C*, red), but State C* is less stable, as indicated by the asymmetric equilibrium arrows. (**B**) Overlay of the *G*-*V* relationship predicted by our simplified kinetic model when [D] = 0 and the hERG *G-V* (reproduced from Furutani *et al*., 2019). Red curve illustrates the activation-*V* relationship of blocked channels. (**C**) Simulated time courses of State O occupancy during a voltage protocol used to elicit facilitation. Inset: curves normalized to the maximal value of the drug-free simulation. (**D**) *G*-*V* curves from peak simulated tail currents with [D] = *K*_D_, with (blue) or without (green) a 4 s pre-pulse to +60 mV before each test pulse. *G*-*V* curve with [D] = 0 shown for comparison. Peak tail open probability normalized by peak tail open probability following +40 mV test pulse with [D] = 0. (**E**) State-to-state flux, probability per second, over time during a facilitated test pulse for unblocking (State O* to State O, dotted curve) and closing (State O to State C, solid curve). (**F**) Occupancies of the closed (State C) and closed-blocked (State C*) states over time. (**G**) Predicted *K*_D_ as a function of voltage.

**Figure 3.**
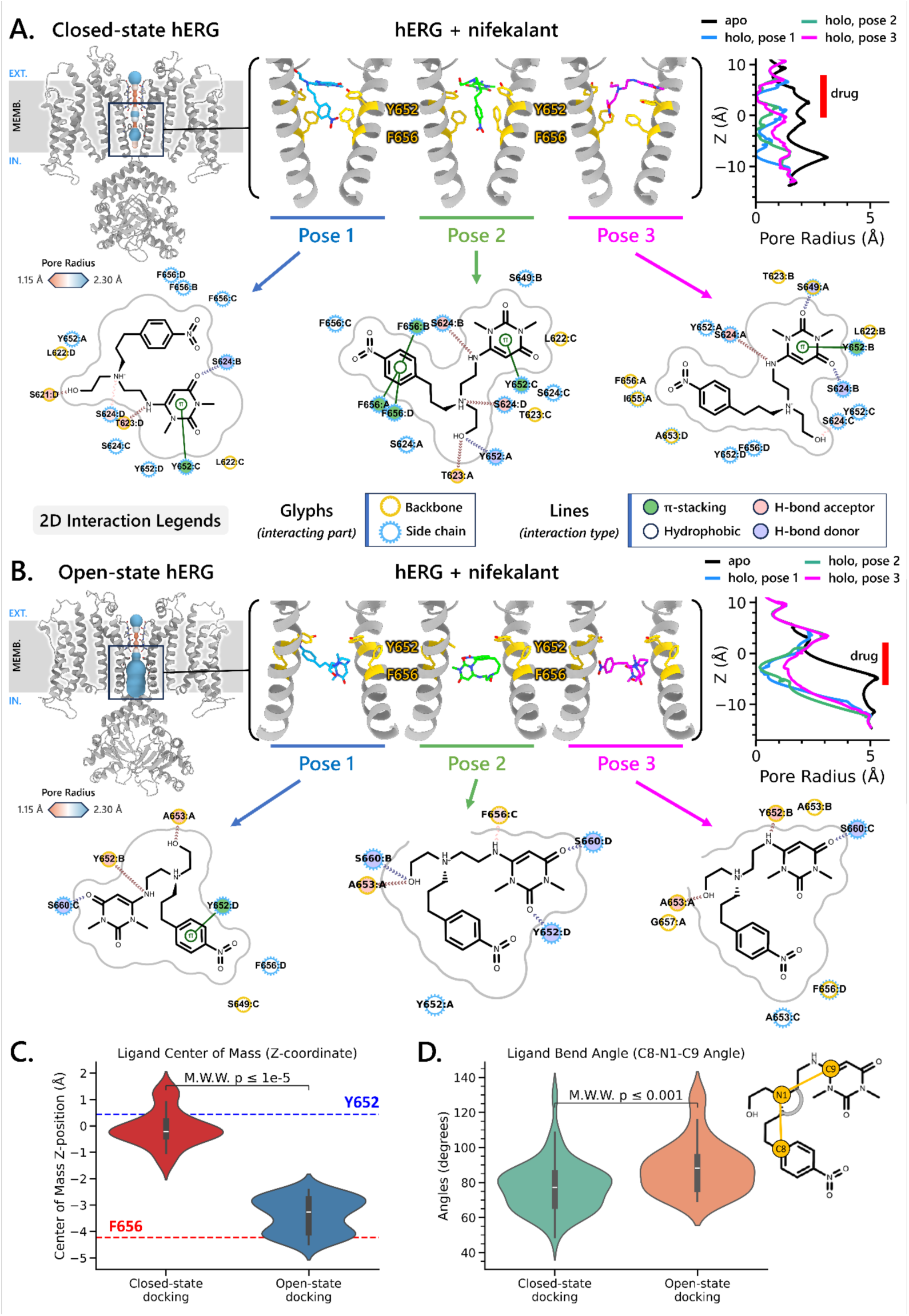
Drug docking simulations reveal distinct nifekalant binding poses in open vs. closed hERG channels. (**A**) closed-state and (**B**) open-state hERG channel models. Three representative binding poses (Pose 1, 2, and 3) were obtained by clustering the top 50 nifekalant docking results with the most favorable binding free energies from a total of 25,000 poses per hERG channel state. For each pose, the pore radius profiles of the hERG channel with the bound drug are shown alongside the *apo* model, at the same vertical level as the visualization directly to the left of the plots. The canonical drug binding residues Y652 and F656 on the pore-lining S6 helices are highlighted in gold. 2D interaction maps are shown to illustrate key contacts between nifekalant and the channel, with interactions visualized for two opposing subunits for clarity. (**C**) Distribution of nifekalant center-of-mass *Z*-position for the top 50 poses (relative to membrane axis) reveals that the drug significantly shifts upward toward the selectivity filter when it binds to the closed-state model compared to open-state model (Mann – Whitney – Wilcoxon U test, M.W.W., *p* ≤ 1e−5). Horizontal dashed lines mark the center-of-mass *Z*-positions of Y652 (blue) and F656 (red) for reference. (**D**) Ligand bend angle, defined as the internal angle formed between the terminal alkyl carbon atoms (C8 and C9) and the central tertiary amine nitrogen (N1), is significantly smaller in the closed-state models (*p* ≤ 0.001), indicating a more compact ligand conformation upon cavity closure.

**Figure 4:**
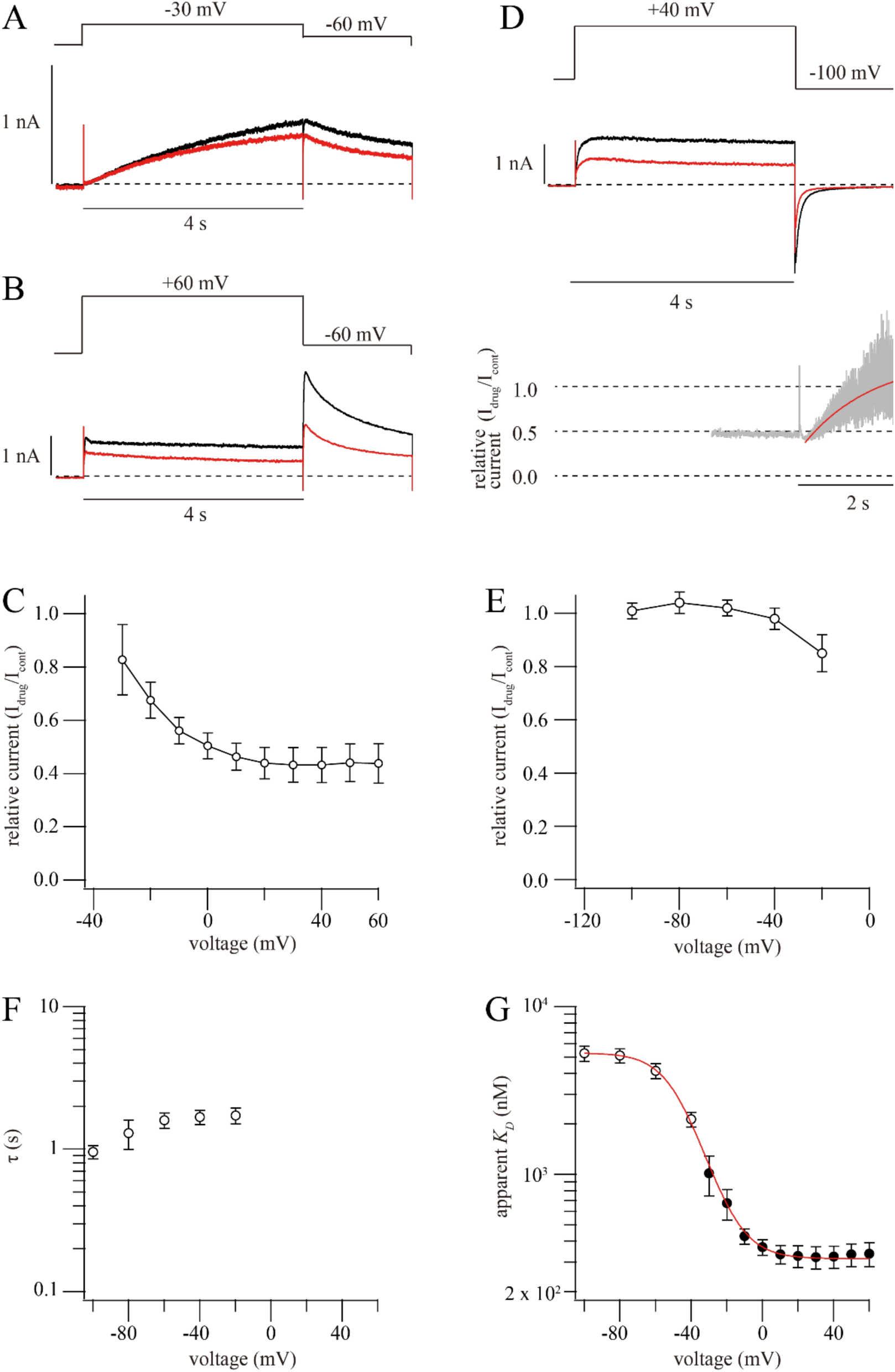
Voltage-dependent affinity suggests that nifekalant prefers activated channels. (**A, B**) Representative whole-cell current recordings showing hERG channel inhibition during test pulses from a holding potential of –90 mV in control conditions (black traces) and after application of 300 nM nifekalant (red traces). (**C**) Relative steady-state current remaining (*I*_drug_/*I*_cont_) as a function of voltage derived from the experiments shown in panels A and B (*n* = 6). (**D**) Upper panel: Representative current traces showing hERG inhibition during test pulses to +40 mV for 4 s, followed by repolarization to –100 mV for 2 s in the control (black) and 300 nM nifekalant (red). Lower panel: Relative steady-state current fit a single-exponential function (red) to tail current ratios (gray) in the presence of 300 nM nifekalant. (**E**) Voltage-dependence of steady-state fractional block (*I*_drug_/*I*_cont_) was quantified from experiments, as shown in panel D (*n* = 5). (**F**) Voltage dependence of the time constants (τ) for nifekalant unbinding. (**G**) Voltage-dependence of nifekalant apparent dissociation constant (*K*_D_). *K*_D_ values were calculated assuming 1:1 binding stoichiometry (open circles: derived from experiments, as in panel D; filled circles: derived from experiments, as in panels A and B; see Methods for details). For highly negative potentials (–60, –80, and –100 mV), 3,000 nM nifekalant was used (*n* = 5) because of its reduced affinity, while 300 nM was used for other potentials (*n* = 5-6). The red curve represents the Boltzmann function. All data are shown as mean ± SEM.

**Figure 5:**
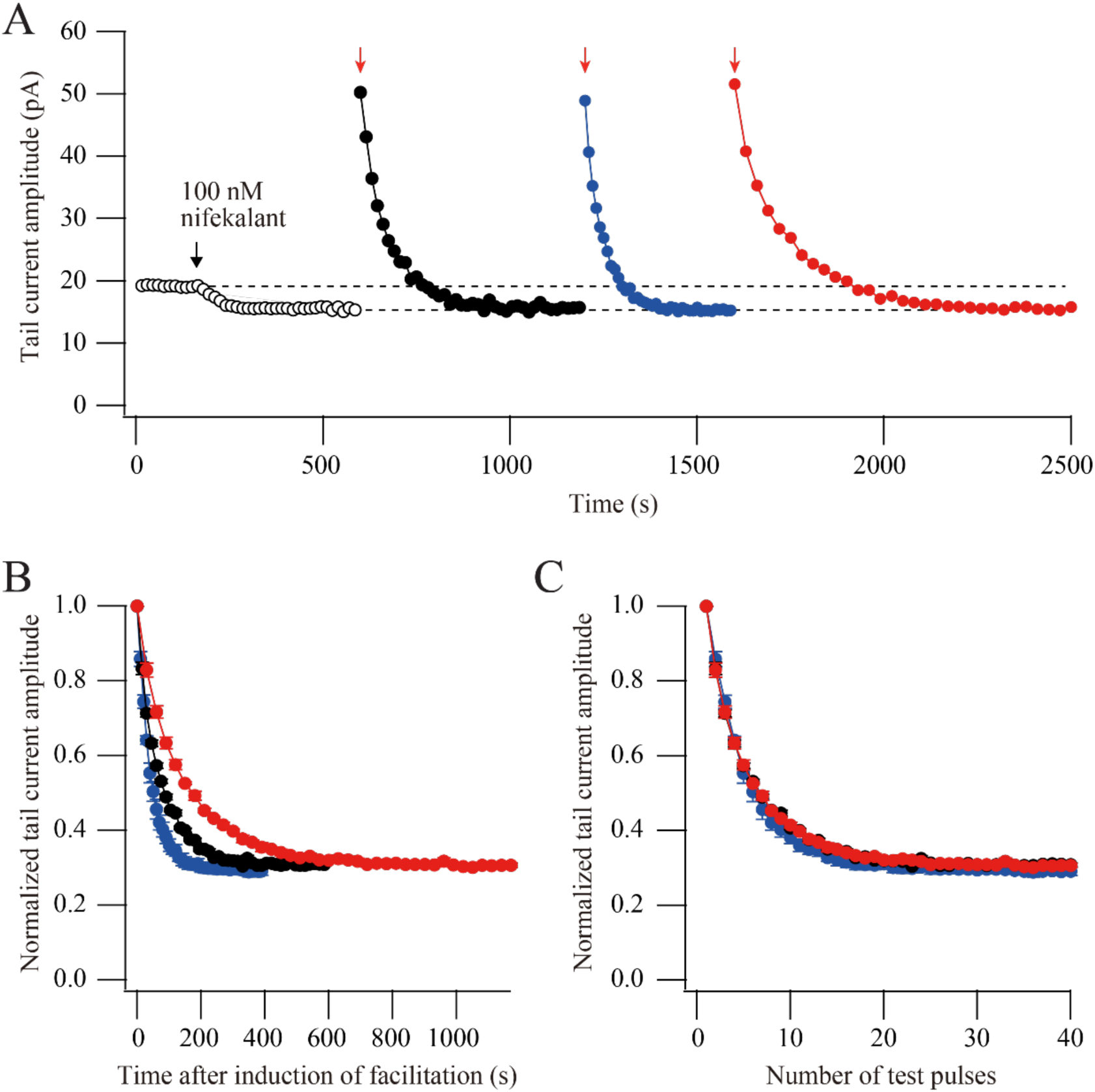
Facilitation decay is use-dependent. (**A**) Representative current recordings demonstrating decay of facilitation during repeated test pulses in the presence of 100 nM nifekalant. Facilitation was induced by 4 s prepulses to +60 mV (indicated by red arrows). Following facilitation induction, test pulses of –40 mV (4 s duration) were applied at intervals of 10, 15, or 30 s. (**B, C**) Time course of the decay of facilitation versus time after prepulse (**B**) and number of test pulses (**C**). Facilitation decayed significantly faster at 10 s intervals than at 30 s intervals (10 s vs 15 s, \**p* < 0.0002; 15 s vs 30 s, \**p* < 0.0001, 10 s vs 30 s, \**p* < 0.0001; with one-way ANOVA followed by Tukey’s post hoc test). No significant differences in decay time constants were observed when the data were normalized to pulse number rather than time (10 s vs 15 s, \**p* = 0.9579; 15 s vs 30 s, \**p* = 0.9752, 10 s vs 30 s, \**p* = 0.9989; with one-way ANOVA followed by Tukey’s post hoc test), indicating that facilitation decay depends primarily on channel activation frequency rather than absolute time. Data are means ± SEM (*n* = 4).

For the detailed drug-*hERG* model, we modified the Adeniran et al. reduced *hERG* channel model (Adeniran *et al*., 2011) by adding drug-bound states and incorporating the proposed drug effects on *hERG* gating kinetics. We did not attempt to reparameterize the Adeniran model to match the kinetics of hERG1a in HEK cells, but rate constants were adjusted from the original 37° C parameterization to 25° C based on the Q_10_ factors listed in Vandenberg et al., 2006 (Vandenberg et al., 2006). As Figure 6A indicates, we restricted drug binding and unbinding to *hERG* channels in the open (𝑂 and 𝑂*) or inactivated states (𝐼 and 𝐼*). We made the simplifying assumption that all activation rate constants of drug-bound channels are altered by the same scaling factor, 𝑓_𝛼_, and all deactivation rate constants are altered by a scaling factor, 𝑓_𝛽_. Furthermore, we assumed that drug binding does not alter the rate constants of inactivation and recovery from inactivation. The full set of differential equations for our detailed drug-*hERG* interaction model are

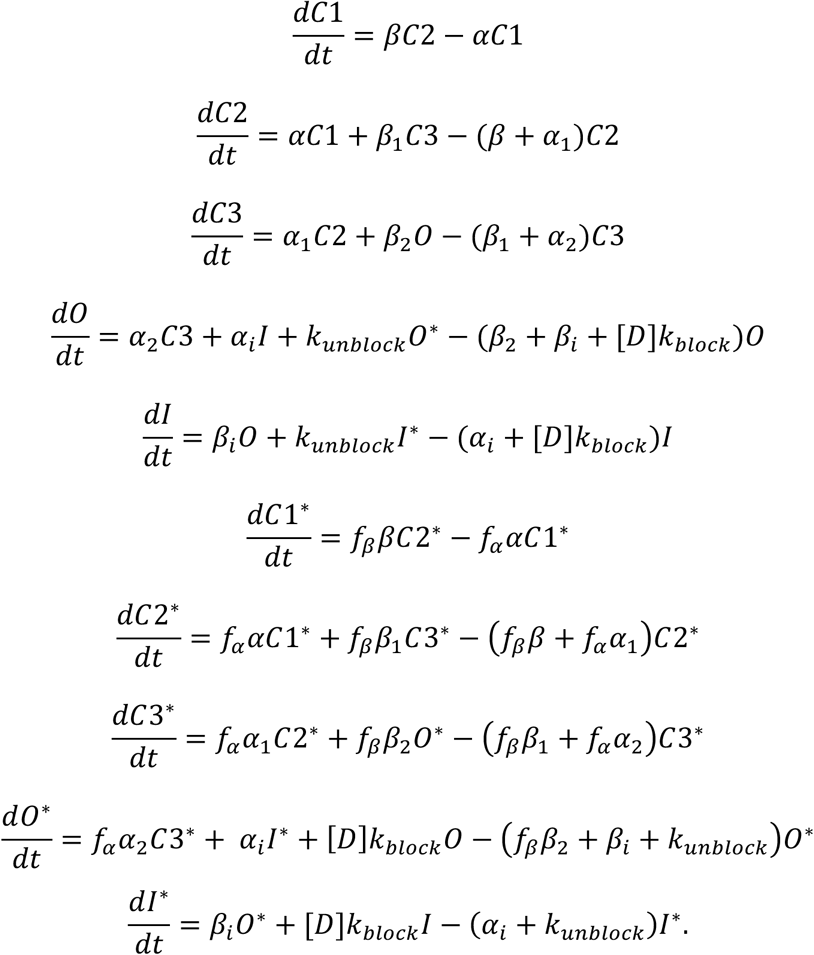

**Figure 6.**
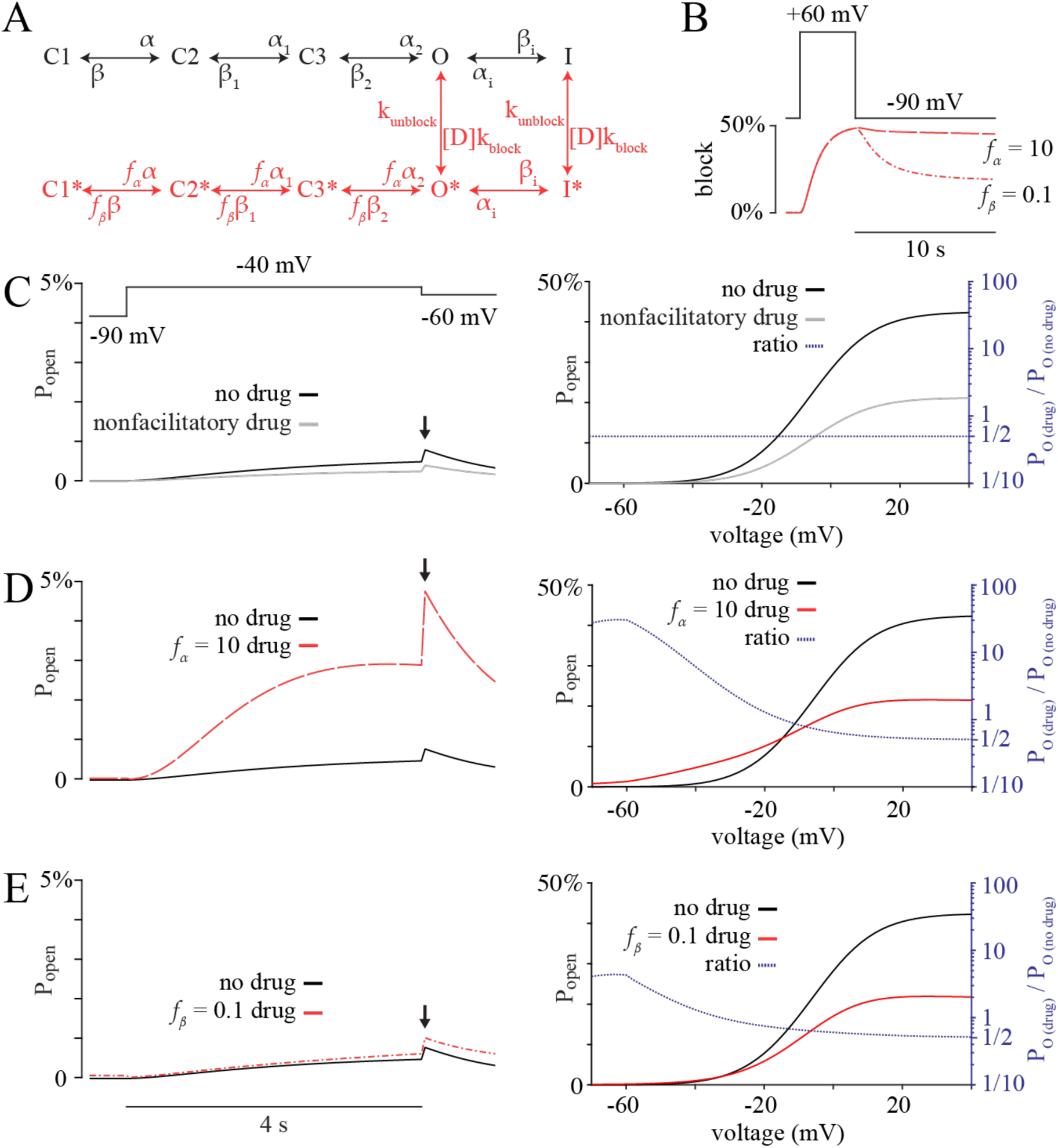
Facilitation in a hERG gating model of agonism-while-blocking. (**A**) blocked states of hERG (red, with asterisks) were added to the hERG gating model developed by Adeniran *et al*. (black) with gating modulation implemented via the factors *f_α_* and *f_β_*. (**B**) Simulated percent block (summed occupancies of states: C1*, C2*, C3*, O*, and I*) during a 4 s pre-pulse to +60 mV and subsequent return to holding potential. (**C-E**) At left, simulated open probability (occupancy of state O) versus time during a –40 mV test-pulse occurring 10 s after a 4 s pre-pulse to +60 mV. Open probability from simulations with [D] = 100 nM (gray and red traces) are overlayed on traces from drug-free simulations (black). Arrows indicate points used for peak open probability quantification. At right, peak open probability during -60 mV step versus test-pulse voltage. Overlaid are ratios of the two peak open probability-voltage curves (blue, right axis), illustrating the degree of facilitation or block as a function of voltage.

Calculations, simulations, analysis, and plotting of both the minimal and detailed drug–hERG channel models were performed using MATLAB R2023b software. For simulations, differential equations were solved using the MATLAB function ode15s, which is a variable time-step solver for stiff differential equations.

## Results

### Agonism-while-blocking can produce facilitation in a simplified kinetic model

We hypothesized that facilitatory blockers bias the S6 gate toward an open conformation while they block, and that facilitation arises from unblock of channels that would otherwise be closed. To assess whether such an agonism-while-blocking mechanism is plausible, we constructed a minimal rate-theory Markov model. The minimal model channel transitions only between a closed state (C) and an open state (O), with corresponding closed/blocked (C*) and open/blocked (O*) states when the drug is bound (Figure 2A). Inactivation was omitted because it is not required for facilitation of hERG by nifekalant (Kushida *et al*., 2002; Hosaka *et al*., 2007). Since nifekalant can only access its blocking site when channels are voltage-activated (Kamiya *et al*., 2006), block and unblock were only permitted from the open state. Voltage-dependent rates of opening and closure for unblocked channels were fitted to the hERG activation kinetics and steady-state open probability (see Methods). Blocking and unblocking rates (*k*_block_ and *k*_unblock_) were set to 5x10^-3^ nM^-1^s^-1^ and 5x10^-1^ s^-1^, roughly matching the kinetics of block by nifekalant and its affinity for maximally voltage-activated channels expressed in HEK cells (Furutani *et al*., 2022). To implement agonism-while-blocking, we scaled the activation rate of blocked channels by a factor *f*_α_ = 40.8 to shift activation *V*_1/2_ by –30mV (Figure 2B), which is similar to the shift observed experimentally during facilitation by nifekalant (Figure 1B).

We then simulated a voltage protocol known to elicit hERG facilitation in the presence of nifekalant. The protocol began at a holding potential of –90 mV and comprised 4 steps. Step 1 was a 4 s pre-pulse to +60 mV, which induces facilitation of hERG by nifekalant *in vitro* (Figure 1). Step 2 returned the membrane potential to the holding potential for 10 s. Step 3 was a 4 s test-pulse to –40 mV, a voltage at which facilitation is robust. Step 4 to –60 mV was for analysis of peak tail current. As a baseline, we simulated this voltage protocol without blocker (Figure 2C). When the drug concentration was set to 100 nM, agonism-while-blocking produced enhanced conduction during the test pulse, whereas setting *f*_α_ = 1 (eliminating agonism) abolished facilitation. Conductance-voltage analysis demonstrated that this enhanced conduction is restricted to test pulses with voltages below the activation *V*_1/2_ and required the activating pre-pulse (Figure 2D), thereby meeting the defining criteria for facilitation. Facilitation in this minimal model suggests that facilitation of hERG conductance may also have a similarly straightforward underlying mechanism.

To understand the mechanism underlying facilitation in the minimal model, we examined state transitions. Importantly, agonism-while-blocking creates state-dependent drug affinity by destabilizing the closed-blocked state. Specifically, by setting *f*_α_ = 40.8, the closed/blocked state is destabilized and the drug affinity for the closed state is weakened by 40.8-fold relative to the open state. Because membrane voltage controls channel state, agonism-while-blocking induces a dependence of dissociation constant (*K_D_*) on voltage, such that block is greater at voltages which open channels (Figure 2G). This state-dependent block drives the dynamics of facilitation.

During the +60 mV pre-pulse, channels open and accumulate blocker. Upon return to –90 mV holding potential, channels rapidly close and trap blocker inside, creating a population of blocked channels that are primed to open during the –40 mV test step because the channels are agonized while they are blocked (Figure 2C,D,F). During the –40 mV test pulse, blocked channels open, but because the blocker affinity at –40 mV is weaker than it was at +60 mV, channels lose their blocker and close, depleting the population of blocked channels (Figure 2E,F). Thus, facilitation occurs when agonized-while-blocked channels become unblocked, revealing open channels.

While this minimal model does not constitute experimental evidence for agonism-while-blocking, it demonstrates that such a mechanism is theoretically sufficient to reproduce the hallmarks of facilitation, and generates three testable predictions: *(1)* The drug always blocks the channel pore when bound, *i.e.* there is no drug-bound conducting state. *(2)* The *K_D_* of the blocker is lower (affinity stronger) at voltages which open channels. *(3)* The blocked state corresponds to the facilitated state. In the following sections, we test these predictions.

### Docking models suggest that nifekalant blocks the central cavity and changes conformation when trapped

The minimal kinetic model predicts that facilitation of hERG could arise from agonism-while-blocking. To assess whether nifekalant could block and agonize, we analyzed *in silico* docking models of nifekalant bound to proposed open and closed conformations of hERG (Ngo *et al*., 2025). We did not consider hERG channel inactivation in these models. Since nifekalant, like other hERG blocking drugs, can be trapped in its hERG blocking site (Kamiya *et al*., 2006), which is proposed to be in the central cavity (Kushida *et al*., 2002; Kamiya *et al*., 2006; Hosaka *et al*., 2007), we first asked whether closed-conformation models of the nifekalant-hERG complex support this trapping behavior. We placed nifekalant in the central cavity of the hERG channel between key binding residues F656 and Y652, and generated 25,000 potential docking poses using Rosetta GALigandDock (Park *et al*., 2021). We then grouped the top 100 structures based on their similarity (see Methods). The top scoring poses from the top three clusters were selected for further analysis. In all cases, nifekalant remained within the central cavity of the closed channel, indicating that the closed channel is spacious enough to accommodate a nifekalant molecule (Figure 3). Furthermore, the S6 helical bundle crossing radius in these closed-bound models was less than 2 Å, too narrow for nifekalant to pass through. This illustrates how nifekalant could become trapped in the central cavity upon channel deactivation.

In the agonism-while-blocking mechanism, nifekalant-bound channels cannot conduct potassium ions. We therefore tested whether potassium ions could potentially pass through nifekalant-bound channels. When docked in the open conformation of hERG, nifekalant fully occludes the ion conduction path, reducing the cavity radius by 70 to 90% (Figure 3B). This leaves open paths with radii of only 0.3 to 1.4 Å, which is too narrow for a hydrated potassium ion with a radius of 3.3 Å to pass through. Representative docked poses show nifekalant interacting with residues from all 4 channel subunits, including Y652 and F656, which are required for block and facilitation by nifekalant (Hosaka *et al*., 2007). Nifekalant spanning the limited space between Y652 and F656 in multiple distinct binding poses suggests a tendency to block ion conduction. Overall, our model suggests that nifekalant-bound channels are always blocked.

The agonism-while-blocking mechanism requires blocked channels to be biased toward opening, so we next asked whether structural features of the hERG-nifekalant complex are consistent with such a state bias, which requires a change in the drug binding conformation upon channel gating. We asked whether nifekalant binds differently to closed hERG channels than open ones. Nifekalant docking poses in open-conformation hERG models are very different from closed-conformation models, with nifekalant moving toward the selectivity filter upon cavity closure (Figure 3C) (Ngo *et al*., 2025). When docked in the open channel, nifekalant sits between the S6 helical bundle crossing point, residues L666, and the sidechains of Y652, which protrude into the cavity (Figure 3B,C). When docked in the closed channel, nifekalant sits farther from L666, extending from F656 past Y652 towards the bottom of the selectivity filter residues S624 (Figure 3A,C). Different open-conformation and closed-conformation docking sites can be attributed to changes in the structure of the central cavity as the channel opens and closes. When the hERG channel pore closes, the pore-lining S6 helices straighten at a kink located between F656 and Y652 (Fowler and Sansom, 2013). This causes residues located below Y652 to rotate toward the central axis of the pore, disrupting the open-conformation nifekalant binding site by reducing the pore radius between residues L666 and Y652. This offers a physical explanation for why hERG channel closure may force nifekalant to move toward the selectivity filter. All representative open-conformation binding poses are incompatible with the closed conformation of hERG because they place nifekalant between Y652 and L666. The Supplemental Movie displays a morph illustrating various binding poses which nifekalant may adopt in the open and the closed conformations of hERG. Overall, these models suggest that the drug-channel binding interface changes when the central cavity closes, consistent with conformational state bias.

The equilibrium between open-bound and closed-bound states also depends on the conformation of nifekalant. In the closed hERG cavity, nifekalant adopts a more compact conformation. This is characterized by a bend around the central tertiary amine moiety of the drug, bringing the two terminal aromatic rings together (Figure 3D). This facilitates the docking of nifekalant within the narrower confines of the closed cavity (Supplemental Movie). We speculate that a more compact conformation of nifekalant may act like a compressed spring, forcing the channel to open around it. Overall, comparison of nifekalant docking poses to open and closed hERG channels indicates that if nifekalant is in the central cavity, it blocks K^+^ conduction and changes its binding pose upon channel closure, as expected of a gating modulator.

Lowest-energy docked poses in both open and closed channels show nifekalant interacting with residues Y652 and F656 and sometimes with S649 and A653. We noticed that nifekalant only contacts G657 and S660 in the open channel and only contacts S621-S624 and I655 in the closed channel (Figure 3A,B). Interactions with these residues may make key contributions to the free energy difference between open-blocked and closed-blocked channels. Facilitation is attenuated by mutations G657A, S624A, and S624T and is enhanced by S660A but none of these mutations affect pore block (Hosaka *et al*., 2007). The distinct interactions of nifekalant with open vs. closed channels offer an explanation for the importance of these residues for facilitation.

### Voltage-activation of hERG underlies voltage dependent affinity of nifekalant

If block by nifekalant biases hERG toward the open state, then the principle of coupled equilibria dictates that nifekalant must have a stronger affinity for the open state than the closed state of the channel. Prior studies have revealed voltage dependence of hERG block in response to some stimulus protocols, but not others (Kushida *et al*., 2002; Hosaka *et al*., 2007; Furutani *et al*., 2022). Here, we measured the degree of block by nifekalant over a wider range of voltages than have been reported for wild-type hERG. Our measurements indicated that the degree of block plateaued above +10 mV but diminished at more negative voltages (Figure 4A,B,C). For voltage steps below -30 mV where instantaneous tail currents were too small for accurate block measurements, we analyzed hERG currents as nifekalant block re-equilibrated during channel deactivation (Figure 4D). By fitting these currents with exponential functions, we obtained a measure of steady-state block along with a reasonable time constant of blocker equilibration (Figure 4E,F). Both approaches yield measures of the degree of steady-state block, reflecting the equilibrium between drug binding and unbinding and allowing direct comparison across the full voltage range. We converted degrees of steady-state block to apparent dissociation constants, *K_D_*, according to a 1:1 Langmuir binding isotherm (Figure 4G). The voltage-dependence of these apprarent *K_D_* values formed a Boltzmann sigmoid distribution on a semi-log plot, as expected of a voltage-dependent conformational change that alters affinity. Fitting a Boltzmann sigmoid function to log *K_D_* values revealed a voltage midpoint of –31 ± 5 mV and a slope of 2.4 ± 0.3 e_0_, similar to the voltage dependence of hERG activation during facilitation by nifekalant, which we reported previously: midpoint of –32 ± 0.7 mV, slope of 2.5 ± 0.1 e_0_ (Furutani *et al*., 2019). This fitting returned *K_D_* values of 5310 ± 50 nM for saturation at negative voltages and 320 ± 50 nM for saturation at positive voltages, a 17 ± 3-fold change. The steep voltage-dependent transition between affinities that saturates at extreme voltages is not consistent with voltage-dependent block (Woodhull 1973). Rather, this pattern is indicative of dependence on voltage-dependent conformational change of a binding site, which for nifekalant would be in the central cavity of hERG. We propose that voltage-dependent activation of hERG underlies this affinity difference: that the open conformation of hERG has ∼17-fold better affinity than the closed state, and consequently that nifekalant block biases hERG gating ∼17-fold towards the open/blocked state.

### Nifekalant trapping underlies the facilitated state

Another key prediction of the agonism-while-blocking mechanism is that the blocked state of hERG corresponds to the facilitated state. If this prediction is accurate, then trapping nifekalant in the central cavity of hERG should also trap channels in the facilitated state. To test equivalence of the blocked and facilitated states, we asked whether maintaining channels in the closed state after a facilitating pre-pulse preserves facilitation. Following a pre-pulse that induces facilitation, the level of facilitation decays during repetitive test pulses (Kamiya *et al*., 2006; Hosaka *et al*., 2007; Furutani *et al*., 2011), but it had remained unclear whether this decay of facilitation results from the repeated pulses or merely reflects dependence on elapsed time from the pre-pulse. We therefore varied test pulse frequency following a facilitating pre-pulse (Figure 5A). We found that facilitation decayed more slowly at lower stimulus frequencies (Figure 5B). When normalized current amplitude was plotted against applied test pulses number, exponential fits yielded indistinguishable decay time constants of 4.5 ± 0.1 pulses, 4.5 ± 0.1 pulses, and 4.5 ± 0.1 pulses when interpulse intervals were 10, 15, and 30 s, respectively (Figure 5C). If nifekalant were able to escape its binding site during the interpulse interval at -90 mV, facilitation would decay faster at longer intervals. The fact that decay depends on pulse number rather than elapsed time indicates that nifekalant cannot escape between pulses. These data suggest that nifekalant remains trapped in its facilitating binding site during voltage-dependent channel closure, much like trapping of nifekalant in its blocking site in the central cavity.

To determine whether the kinetics of facilitation decay are consistent with relief of block by nifekalant, we analyzed nifekalant dissociation kinetics. Facilitation decays with a characteristic time constant of ∼18 s (cumulative time at –40 mV, Figure 5C), even though the S6 gates are open, allowing the drug to escape, only during a fraction of this time. This implies that the escape rate when S6 gates are open is faster than the decay rate of facilitation. The escape time constant of 1.7 ± 0.2 s at –40 mV from channels that have not yet closed their activation gates (Figure 4F) is substantially faster than the 18 s time constant during depolarizing steps to –40 mV (Figure 5C). This finding is consistent with nifekalant being trapped in its binding site for facilitation in closed channels. Altogether these results support the hypothesis that the facilitated state and the blocked state are the same.

### Agonism-while-blocking produces facilitation in an established model of hERG gating

To further assess whether the agonism-while-blocking observed with nifekalant could produce facilitation, we implemented agonism-while-blocking in an established Markov model of hERG gating (Adeniran *et al*., 2011). In this model, gating is represented by sequential transition between three closed states (C1, C2, and C3), an open conducting state (O), and an inactivated state (I). To incorporate drug binding effects, we assume that drug-bound hERG channels can occupy the corresponding states: C1*, C2*, C3*, O*, and I* (Figure 6A). As in the minimal model, we forbade drug exchange with closed channels, and we assumed that drug binding did not alter inactivation kinetics. The drug binding and unbinding rates (*k*_block_ and *k*_unblock_) for open or inactivated channels were set as in the minimal model. Agonism-while-blocking was implemented by scaling activation and deactivation rates for drug-bound channels by the factors *f*_α_ and *f*_β_, respectively. We refer to this as our detailed model of agonism-while-blocking in hERG. Using the detailed model, we simulated voltage protocols similar to those used with the minimal model, and peak tail conductances were used for analysis, as is typical for hERG conductance analyses.

We tested whether agonism-while-blocking could produce facilitation in the detailed hERG model. We first implemented agonism-while-blocking by either slowing deactivation tenfold (*f*_β_ = 0.1) or speeding activation tenfold (*f*_α_ = 10). As in the minimal model, the pre-pulse loaded channels with drug, that became trapped when channels closed upon return to the negative holding potential (Figure 6B). No facilitation was observed when block occurred without agonism (*f*_α_ = *f*_β_ = 1; Figure 6C), whereas either slowing deactivation or accelerating activation reproduced the defining features of facilitation (Figure 6D,E). Increasing *f*_α_ produced a larger increase in test pulse open probability than decreasing *f*_β_ by the same factor, resulting, in part, from better trapping after loading (Figure 6B). To explore this difference systematically, we independently varied *f*_β_ and *f*_α_ over a range from 0.01 to 100. Increasing *f*_α_ or decreasing *f*_β_ initially increases test pulse conductance and then decreases it, with *f*_α_ producing a larger maximal effect (Figure 7A). Mapping this parameter space reveals a ridge of maximal facilitation when the ratio of *f*_α_ and *f*_β_ is ∼9 and *f*_β_ > 1 (Figure 7B).

**Figure 7.**
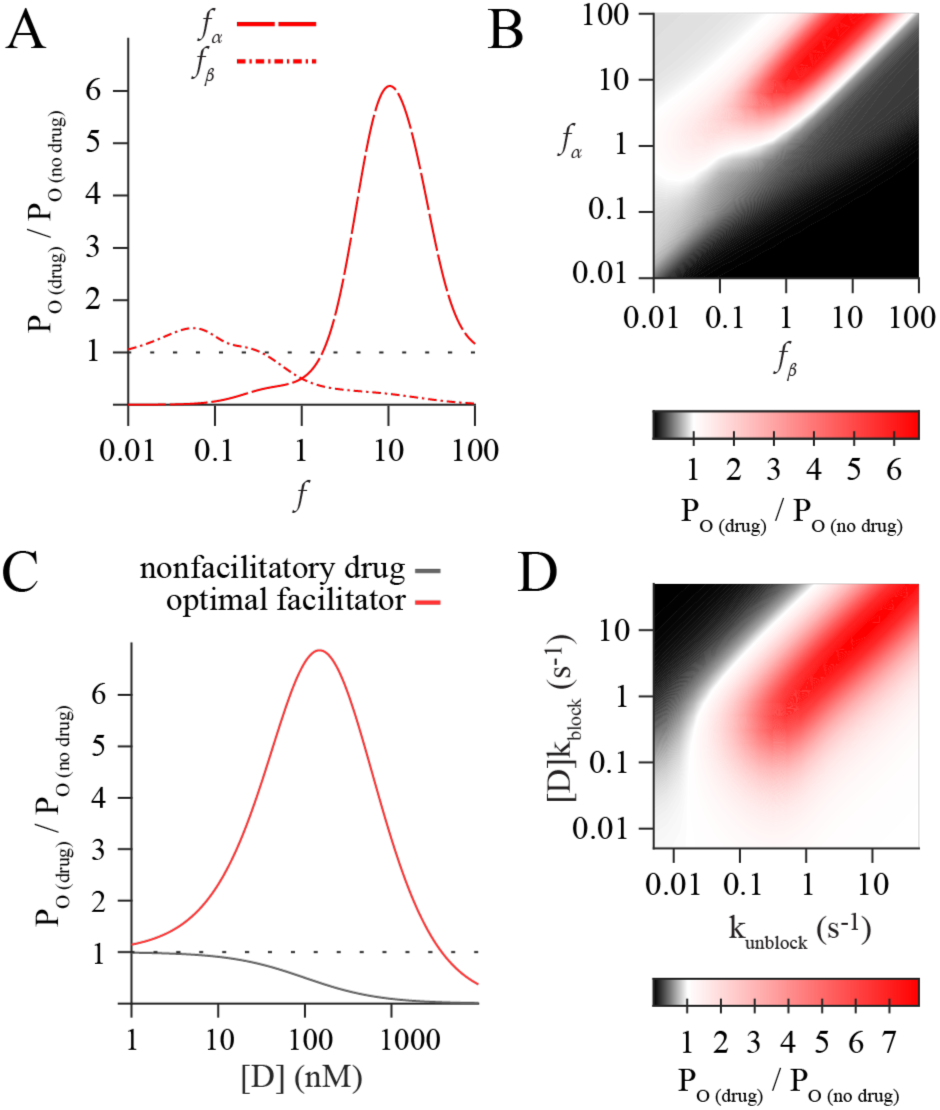
Optimization of a hERG facilitation model. Maximum open probability during the –60 mV tail step (following a test-pulse to –40 mV) in simulations. Open probabilities were normalized to control, drug-free simulation. (**A**) *f_α_* and *f_β_* were individually varied. (**B**) Heatmap of maximum open probability when *f_α_* and *f_β_* are both varied. (**C**) Drug concentration [D] was varied. Optimal facilitator has *f_α_* = 92 and *f_β_* = 10. (**D**) Drug on and off rates were varied for the optimal facilitator.

We next examined how facilitation depends on drug concentration by varying drug levels from 1 to 10,000 nM for a non-facilitatory blocker (*f*_α_ = *f*_β_ = 1) and for an ‘optimal facilitator’ with *f*_α_ = 92 and *f*_β_ = 10 (Figure 7C). Our model reproduced the expected concentration-response relationships for both types of blockers: the non-facilitatory blocker reduced test pulse conductance according to a single-binding-site Hill curve, whereas the optimal facilitator produced a biphasic response in which facilitation peaked near the open state *K_D_* of 100 nM. We then examined how the kinetics of drug exchange influenced facilitation by systematically varying [D]*k*_block_ and *k*_unblock_ for a facilitatory blocker. The resulting heat map (Figure 7D) shows that facilitation is maximal in a region where [D]*k*_block_ is approximately equal to *k*_unblock_, corresponding to ∼50% block when channels are in open or inactivated states. Consistent with this prediction, the half-maximal effective concentration, EC_50_, of facilitation is typically near the IC_50_ for block by facilitatory drugs (Yamakawa *et al*., 2012). The ridge disappears at *k*_unblock_ values below ∼1 s^-1^, where unblock becomes slower than the 1.8 s^-1^ channel deactivation rate, causing channels to close before they can be unblocked. This constraint highlights a key requirement of the agonism-while-blocking mechanism: channels opened by an agonizing blocker must dwell in the open state long enough after unbinding to contribute to a macroscopic conductance. Overall, our model indicates that facilitation is strongest at concentrations that block ∼50% of channels.

Lastly, we investigated the state occupancies in simulations of the detailed model to determine whether facilitation arises through a mechanism similar to that observed in the minimal model (Figure 2), despite the detailed model having multiple closed states and an inactivated state. Simulations with the optimal facilitator showed that occupancy of the inactivated state was proportional to the open state during facilitation (Figure 8A,B). As in the minimal model, facilitation was accompanied by progressive loss of bound blocker during the test pulse (Figure 8C) and accumulation of channels in unblocked closed states followed activation (Figure 8D). Examination of the state-to-state fluxes revealed the overall flow during facilitation: channels transition from blocked closed states to blocked open states, then to unblocked open states, and finally to unblocked closed states (Figure 8E). These results demonstrate that the detailed model reproduces the same core mechanism identified in the minimal model, showing how a blocker that promotes channel activation can indeed produce facilitation of hERG conductance through an agonism-while-blocking mechanism.

**Figure 8.**
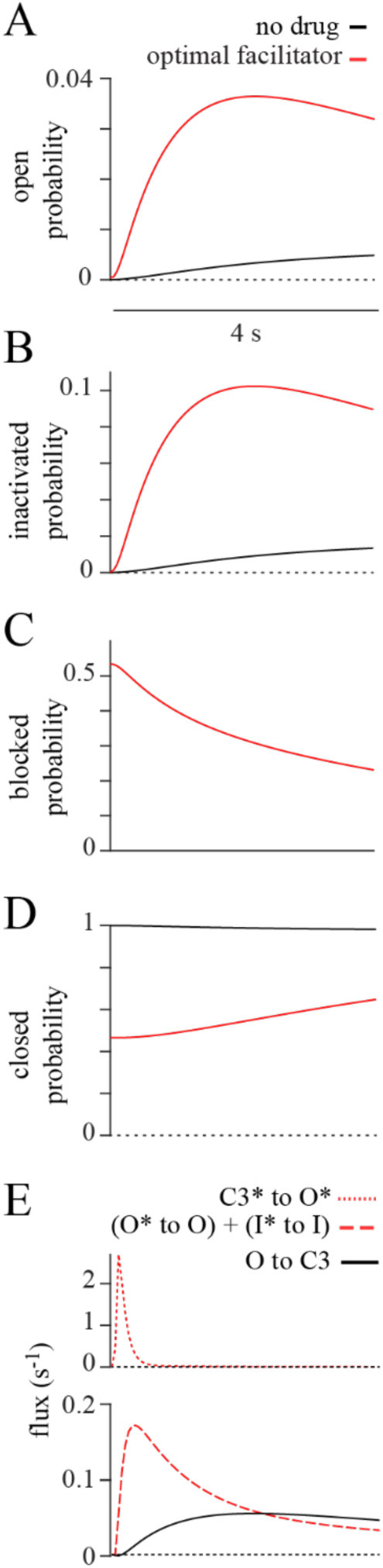
State transitions underlying facilitation: (**A**) Simulated hERG state O occupancy (open probability) during a step to –40 mV with a maximally facilitatory blocker: *f_α_* = 92, *f_β_* = 10, [D] = 100 nM = *K*_D_. (**B**) Occupancy of state I. (**C**) Summed occupancy of blocked states C1* + C2* + C3* + O* + I*. (**D**) Summed occupancy of unblocked closed states C1 + C2 + C3. (**E**) State-to-state flux.

## Discussion

We propose that facilitation results from agonism-while-blocking: blockers promote channel opening and then dissociate to reveal conductive channels. Agonism-while-blocking is related to classical foot-in-the-door blockade of voltage-gated channels by quaternary ammonium ions which interfere with channel closing (Yeh and Narahashi, 1977). Agonism-while-blocking is also conceptually related to the inactivation-interference mechanism by which blocker 4-aminopyridine increases macroscopic conductance of Kv2.1-containing channels (Fernández-Mariño *et al*., 2023). Facilitation by agonism-while-blocking is implicitly limited to pore blockers that dissociate on the time scale of channel closure or faster. We therefore do not expect facilitation to be induced by blockers with very slow dissociation rates *e.g.* dofetilide (Tsujimae *et al*., 2004), bepridil, domperidone, and E-4031 (Stork *et al*., 2007). We conclude that agonism-while-blocking underlies facilitation and that facilitation can only occur with blockers that dissociate with kinetics similar to or faster than hERG deactivation.

### Limitations

It is unclear to what degree the atomistic details of modeled nifekalant-hERG interactions are physiologically predictive. The structural modeling here indicates that the conformation of nifekalant within the hERG cavity blocks K^+^ passage. However, there could be an unexpected path for K^+^ to bypass the blocker. Different affinities to open vs. closed hERG channels are required by the agonism-while-blocking mechanism, and modeling indicates that the nifekalant binding pose needs to change between open and closed channels. However, molecular docking calculations did not reveal significantly different affinities between the distinctive open and closed binding poses of nifekalant (Ngo *et al*., 2025).

We did not attempt to find a precise quantitative fit of our detailed drug-hERG model to the data presented here, as our experimental conditions are distinct from model parameterization conditions (Adeniran *et al*., 2011). Our experiments are limited to nifekalant and heterologously-expressed hERG1a homomers in near-physiological solutions at room temperature. The kinetic model also has a simplifying assumption that drug binding affects all closed states equally, as we lack a basis for distinguishing between the affinities of closed states. Our estimates of nifekalant affinity at various voltages are likely affected by knock-on (Hodgkin and Keynes, 1955) of permeant ions with nifekalant, which we did not account for. We note, however, that any knock-on effects are likely small as nifekalant affinity is only modestly changed when hERG D540K mutant channels are opened to inward currents at extreme negative voltages (Furutani *et al*., 2022).

Further work will be needed to establish whether other facilitatory blockers share the same mechanism and how facilitation is affected by *in vivo* differences such as expression of the hERG 1b isoform and changes in biochemical regulation. Nonetheless, our simulations suggest that facilitation is robust across a range of blocker kinetics and affinities (Figure 7D) and so is likely to occur with physiological hERG gating kinetics.

### Applicability of the agonism-while-blocking model to other facilitatory blockers

To date, at least 15 compounds have been shown to exhibit facilitation as defined here (Jiang *et al*., 1999; Furutani *et al*., 2011; Yamakawa *et al*., 2012). Several other hERG blockers can act as agonists during certain voltage protocols (Table 1), raising the question of whether they share a common mechanism of action. Kinetic and mutational data support a central cavity blocking site for nifekalant and other blockers including almokalant (Carmeliet, 1993), amiodarone (Kiehn *et al*., 1999), chlorpheniramine (Hong and Jo, 2009), fluoxetine (Thomas *et al*., 2002), haloperidol (Stork *et al*., 2007; Suessbrich *et al*., 1997), imipramine (Teschemacher *et al*., 1999), promethazine (Jo *et al*., 2009), quinidine (Sănchez-Chapula *et al*., 2003), and verapamil (Zhang *et al*., 1999). Azimilide is unique among known facilitatory blockers in that it blocks only from the extracellular side of the membrane (Jiang *et al*., 1999) and so must act by a different mechanism.

**Table 1.** Compounds exhibiting the kinetic properties of facilitation. Citations: ^1^(Abi-Gerges et al., 2011); ^2^(Amos et al., 2003); ^3^(Busch et al., 1998); ^4^(Carmeliet, 1993); ^5^(Chouabe et al., 1998); ^6^(Duan et al., 2007); ^7^(Dupuis, Klaerke and Olesen, 2005); ^8^(Furutani et al., 2022); _9_(Furutani et al., 2011); ^10^(Hong and Jo, 2009); ^11^(Hosaka et al., 2007); ^12^(Jiang et al., 1999); ^13^(Jo et al., 2009); ^14^(Kamiya et al., 2006); ^15^(Kauthale et al., 2015); ^16^(Kawakami et al., 2006); ^17^(Kiehn et al., 1999); ^18^(Kushida et al., 2002); ^19^(Sănchez-Chapula et al., 2003); ^20^(Stork et al., 2007); ^21^(Suessbrich et al., 1997); ^22^(Teschemacher et al., 1999); ^23^(Thomas et al., 2002); ^24^(Tsujimae et al., 2004); ^25^(Waldegger et al., 1999); ^26^(Walker et al., 2000); ^27^(Yamakawa et al., 2012); ^28^(Zhang et al., 1999); ^29^(Zhang et al., 2016).

| Compound<br>(PubChem CID) | Prepulse<br>$\Delta V_{1/2}^*$<br>(mV) | State<br>preference | Voltage<br>dependent<br>affinity** | Use<br>dependent<br>affinity | Activation<br>required<br>for block | Trapping | Miscellaneous |
| --- | --- | --- | --- | --- | --- | --- | --- |
| Almokalant<br>(3033962) |  | O/I<sup>2,4</sup> | Yes<sup>2,4</sup> | Yes<sup>4</sup> | Yes<sup>4</sup> | Yes<sup>4</sup> | Lower affinity<br>for hERG1b-containing<br>channels<sup>1</sup>. |
| Amiodarone<br>(2157) | -29.9<sup>9</sup> | O/I<sup>17</sup> | Yes<sup>17</sup> | Yes<sup>17,20</sup> | Yes<sup>17,20,29</sup> |  | Higher affinity<br>at higher<br>temperature<sup>15</sup>. |
| Azimilide<br>(9571004) |  | O/I<sup>3,12,26</sup> | Yes<sup>26</sup> | Yes<sup>3</sup> | Yes<sup>3</sup> |  | Blocks<br>externally<sup>12</sup> and<br>facilitation<br>requires<br>inactivation<sup>12,26</sup>. |
| Carvedilol<br>(2585) | -16.0<sup>27</sup> |  |  | No<sup>16</sup> |  |  |  |
| Chlorpheniramine<br>(2725) | -12.9<sup>27</sup> | O/I<sup>10</sup> | Yes<sup>10</sup> |  | Yes<sup>10</sup> |  | Higher affinity<br>for hERG1b-containing<br>channels<sup>1</sup>. |
| Fluoxetine<br>(3386) | -20.9<sup>27</sup> | O<sup>23</sup> | Yes<sup>23</sup> | No<sup>23</sup> | Yes<sup>23</sup> | | Higher affinity<br>for hERG1b-containing<br>channels<sup>1</sup>.<br>Negatively<br>shifts the<br>inactivation<br>$V_{1/2}^{23}$. |
| Haloperidol<br>(3559) | -20.5<sup>27</sup> | I<sup>20,21</sup> | Yes<sup>20</sup> | Yes<sup>20,21</sup> | Yes<sup>20,21</sup> |  | Increases the<br>rate and degree<br>of<br>inactivation<sup>21</sup>. |
| Imipramine<br>(3696) | -10.3<sup>27</sup> | O/I<sup>22</sup> | Yes<sup>22</sup> |  | Yes<sup>22</sup> |  | Higher affinity<br>for hERG1b-containing<br>channels<sup>1</sup>. |
| Metoprolol<br>(4171) | -14.7<sup>27</sup> |  |  | No<sup>16</sup> |  |  |  |
| Nifekalant<br>(4486) | -27.2<sup>9</sup> | O<sup>8,11,14,18</sup> | Yes<sup>8,14,18</sup> | Yes<sup>18</sup> | Yes<sup>8,18</sup> | Yes<sup>14</sup> | Slows channel<br>deactivation,<br>accelerates<br>inactivation and<br>recovery, and<br>negatively<br>shifts the<br>inactivation<br>$V_{1/2}^{18}$. |
| Nortriptyline<br>(4543) | -14.1<sup>27</sup> |  |  |  |  |  |  |
| Promethazine<br>(4927) | -23.6<sup>27</sup> | O/I<sup>13</sup> | Yes<sup>13</sup> |  | Yes<sup>13</sup> |  |  |
| Propranolol<br>(4946) | -12.3<sup>27</sup> |  | Yes<sup>7</sup> | No<sup>16</sup> |  |  | KCNE2<br>coexpression increases affinity<sup>7</sup>. |
| Quinidine (441074) | -21.7<sup>9</sup> | O/I<sup>19,24</sup> | Yes<sup>19,24</sup> | No<sup>24</sup> | Yes<sup>19,24</sup> |  | Slows tail current deactivation<sup>24</sup>. |
| Verapamil (2520) | -13.0<sup>27</sup> | O/I<sup>5,6,25,28</sup> | Yes<sup>5,6</sup> | Yes<sup>28</sup> | Yes<sup>5,6,28</sup> | Yes<sup>28</sup> | Slows channel deactivation<sup>28</sup> and negatively shifts the inactivation $V_{1/2}$<sup>5,6</sup>. |
| Zenidolol (104824) | -13.4<sup>27</sup> |  |  | No<sup>16</sup> |  |  |  |

\* The change in the half-maximal activation voltage of facilitated channels, as assessed by double Boltzmann fits to activation voltage relations in the presence of facilitators. Adapted from (Furutani et al., 2011) and (Yamakawa et al., 2012).
\*\* In most articles cited here, assays do not distinguish between intrinsic voltage dependence of block and the effects of channel gating. The voltage dependence of nifekalant (Kushida et al., 2002), and almokalant (Carmeliet, 1993) have been shown to depend on channel gating.

Trapping can hold channels in the facilitated state – an apparent advantage for enhancing channel conductance. Trapping in closed channels is evident for nifekalant (Kamiya *et al*., 2006), almokalant (Carmeliet, 1993), and verapamil (Zhang *et al*., 1999). However, our data do not address whether trapping is required for agonism-while-blocking.

Evidence for preferential binding to open cavities over closed cavities has been observed for most known facilitatory blockers. For some blockers, this is seen as an increase in percent block at more positive voltages. Block increases with increased voltage for chlorpheniramine (Hong and Jo, 2009), imipramine (Teschemacher *et al*., 1999), propranolol (Dupuis, Klaerke and Olesen, 2005), and promethazine (Jo *et al*., 2009). Some blockers unbind from their blocking or facilitating sites while membranes are held at sufficiently negative voltages. This behavior indicates weaker affinity for closed than for open channels. Compounds with this behavior are: nifekalant (Hosaka *et al*., 2007; Kushida *et al*., 2002); almokalant (Carmeliet, 1993); amiodarone (Kiehn *et al*., 1999; Lo and Kuo, 2019); haloperidol (Stork *et al*., 2007); quinidine (Sănchez-Chapula *et al*., 2003; Tsujimae *et al*., 2004); and verapamil (Zhang *et al*., 1999). We note that several facilitatory blockers, including almokalant, have exhibited their hallmark agonism without explicit application of a prepulse (Table 1).

More generally, the agonism-while-blocking model predicts that perturbations to channel gating kinetics will influence facilitation. Consistent with this expectation, some facilitatory blockers have different affinities for hERG1b-containing channels compared to hERG1a homotetramers (Abi-Gerges *et al*., 2011), which differ in gating kinetics. Whether such affinity differences translate to differences in facilitation remains to be tested.

### Implications for cardiac pharmacology

A remarkable number of facilitatory hERG blockers remain clinically approved. It is possible that facilitatory blockers are safer because they enhance hERG conduction during the downstroke of the cardiac action potential (Furutani *et al*., 2019). It is also possible that agonism-while-blocking undergirds an anomalous safety factor for some hERG blockers. Presently, hERG inhibition is grounds for rejection of drug candidates (Leishman, Abernathy and Wang, 2020), which may potentially eliminate safe and efficacious leads. Our results provide a mechanistic hypothesis for why some hERG-blocking drugs may be safer and rescue promising drug candidates.

## Acknowledgments

We are grateful to Dr. Mark T. Keating and Dr. Michael C. Sanguinetti (University of Utah) for providing us with the *hERG* clone and Dr. Craig T. January (University of Wisconsin) for providing us with HEK293 cell lines stably expressing *hERG*. This study was supported by US National Institutes of Health grants U01HL126273, R01HL128537, R01HL174001, and R01HL168874, UC Davis T32 training program in Basic and Translational Cardiovascular Science (T32HL086350) as well as the Center for Precision Medicine and Data Sciences at UC Davis Health, a Dean’s Award and Bridge Funding Award from the Department of Physiology and Membrane Biology, University of California, Davis, Oracle for Research fellowship, and the Scientific Research (C) 21K06812 and 24K10045 from the Ministry of Education, Science, Sports and Culture of Japan, the Japan Society for the Promotion of Science. Computer allocations were provided through US National Science Foundation Advanced Cyberinfrastructure Coordination Ecosystem: Services & Support (ACCESS) grants BIO250260 and MCB170095. OpenEye’s free academic license was provided by Cadence Molecular Sciences.

## Data and code availability

Code and data for the mathematical simulations in this study will be made public upon publication at the following site: https://github.com/ssdocken88/hERG_blocker_conductance_increase

